# Influenza virus fusion at the plasma membrane is restricted by liquid-ordered membrane components

**DOI:** 10.1101/2025.03.28.645900

**Authors:** Steinar Mannsverk, Ana M. Villamil Giraldo, Peter M. Kasson

## Abstract

Many enveloped viruses enter cells via fusion with the endosomal membrane, raising the question whether entry through the endosomal route confers a fitness advantage over fusion directly at the plasma membrane. We found that influenza A virus fusion at the plasma membrane of A549 cells resulted in a 7.6-fold reduction in productive cell infection, compared to infection through the physiological endosomal route. We hypothesized that this was partially explained by restrictive and permissive membrane factors at the plasma and endosomal membrane, respectively. To test this, we developed a single-viral content mixing assay with plasma membrane vesicles (PMVs), where a fluorescent content marker was loaded into purified PMVs at a quenched concentration, through freeze-thawing. We show that influenza fusion with the plasma membrane is 6-fold less efficient than with liposomes containing the endosome-enriched phospholipid BMP. Incorporating BMP into the PMVs with pre-loaded methyl-α-cyclodextrin (MαCD) did not rescue full fusion, suggesting that the lack of BMP in the plasma membrane is not sufficient to explain decreased fusion with the plasma membrane. Depletion of cholesterol from PMVs enhanced lipid mixing rates and reduced membrane order. Lastly, modest depletion of cholesterol from A549 lung epithelial cells rescued influenza infection through fusion at the plasma membrane by 2.4-fold. We propose that for the endosome-adapted influenza virus, fusion directly at the plasma membrane is restricted by the liquid-ordered-like nature of the plasma membrane, raising the question whether the plasma membrane is broadly less permissive to viral entry than endosomes.

**Importance:** Influenza virus normally enters cells via endosomes, yet it has been known for decades that acidic conditions can force it to enter via the plasma membrane. Here, we show that the plasma membrane route yields 9-fold worse infection of cells and, surprisingly, most of this effect can be attributed to the plasma membrane being less permissive to viral membrane fusion. This difference between plasma membrane and endosomes cannot be simply abolished by adding a key endosomal lipid that can boost fusion in synthetic membranes. The outer surface of the plasma membrane is known to be highly ordered, and disrupting this order does increase fusion as well as boost the ability of influenza virus to infect cells. We therefore conclude that the complex and ordered composition of cell membranes helps control when and where influenza virus can enter and infect.

## Introduction

After binding to host cells, many enveloped viruses are internalized by endocytosis. This is followed by viral membrane fusion with the endosomal or endolysosomal membrane, resulting in release of the viral genome into the host cell cytoplasm (1). However, some enveloped viruses fuse directly with the plasma membrane after binding, bypassing the endosomal pathway. These include paramyxoviruses, many retroviruses and, under some circumstances, SARS-CoV-2 (2). Fusion directly at the plasma membrane is typically associated with the trigger for fusion being a co-receptor or protease present at the plasma membrane, whereas endosome-fusing viruses are typically triggered by receptors, proteases, or the acidic pH within endosomes and lysosomes (2).

It has been suggested that viral entry via the endocytic route confers a fitness advantage over fusion directly at the plasma membrane. Some postulated mechanisms for this advantage include hijacking the intracellular transport machinery to reach the site of replication or immune evasion by internalization of immunogenic viral components (1). It is also possible that the endosomal membrane is intrinsically more permissive for fusion of certain viruses than the plasma membrane. Supporting this idea, recent studies have shown that the endosome-enriched phospholipid bis(monoacylglycero)phosphate (BMP) promotes fusion of multiple viruses (3–8).

Endosomal and plasma membranes have very different physical properties, reflecting their different physiological roles. Specifically, the plasma membrane outer leaflet is enriched in sphingomyelin (SM) and cholesterol, resulting in lower membrane fluidity, tighter lipid packing and a membrane able to resist mechanical stress (9). Unsurprisingly, viruses fusing directly with the plasma membrane appear to leverage its unique physical properties. For instance, the HIV-1 fusion peptide preferentially inserts into the boundary between the liquid-disordered and liquid-ordered domains (enriched in cholesterol, SM and saturated phospholipids) and likely acts there to promote fusion (10).

Here, we hypothesize that the physical properties of the plasma membrane may restrict fusion of endosome-adapted viruses. We used influenza A virus (IAV) as a prototype virus, because it is adapted to fuse with mid-to late-endosomes during cell entry and the sole trigger for fusion is low pH (11). To probe whether the plasma membrane restricts IAV fusion, we first tested whether exogenously triggered IAV fusion at the plasma membrane was less efficient in causing productive infection of a respiratory epithelial cell line than entry via the endosomal route. We then purified plasma membrane vesicles (PMVs) from cells and used them to study single-virus fusion within microfluidic flow cells. We found that IAV fusion with PMVs is markedly less efficient than with liposomes containing BMP. However, adding BMP to PMVs was not sufficient to recapitulate the fusion levels observed in BMP-containing liposomes. Interestingly, cholesterol extraction from PMVs both enhanced lipid mixing kinetics with IAV and reduced overall plasma membrane order. Lastly, extracting cholesterol from lung epithelial cells partially rescued productive infection after triggering IAV fusion at the plasma membrane of cells. Our data establish a membrane-specific effect to the IAV preference for fusion in endosomes. They further suggest that the cholesterol-dependent low fluidity of the plasma membrane outer leaflet may restrict fusion by IAV and potentially other endosome-adapted viruses.

## Results

### IAV forced fusion at the plasma membrane

We first measured the quantitative efficiency of IAV infection when fusion was triggered at the plasma membrane versus physiological fusion in endosomes (**Fig. 1a**). We adapted a well-established assay for forced fusion at the plasma membrane, in A549 human lung epithelial cells (12). X-31 (H3N2) influenza A virus (IAV) was bound to cells at 4°C, followed by either canonical endocytosis and fusion within endosomes (**Fig. 1a I**) or forced fusion at the plasma membrane (FFPM); whereby IAV bound to cells is exposed to a pH 5.0 buffer, triggering fusion (**Fig. 1a II**). Infection efficiency was assessed by anti-IAV nucleoprotein (IAV NP) immunostaining 6 hours post-infection (**Fig. 1b)**. Similar to previous reports (13), treatment with a weak base (here NH_4_Cl) completely inhibited infection (**Fig. 1b**). FFPM partially rescued infection, yielding a 7.6-fold decrease in the percentage of infected cells compared to physiological fusion in endosomes (**Fig. 1c**). This suggests that IAV fusion at the plasma membrane can lead to productive cell infection, albeit less efficiently than through fusion from within endosomes. This difference is likely multifactorial, and prior reports have identified portions of the transport and degradation pathways associated with endosomal transport that facilitate infection (14). Here, we test the hypothesis that some of the difference in infection efficiency between the endosomal and plasma membrane stems directly from their difference in membrane factors.

**Figure 1.**
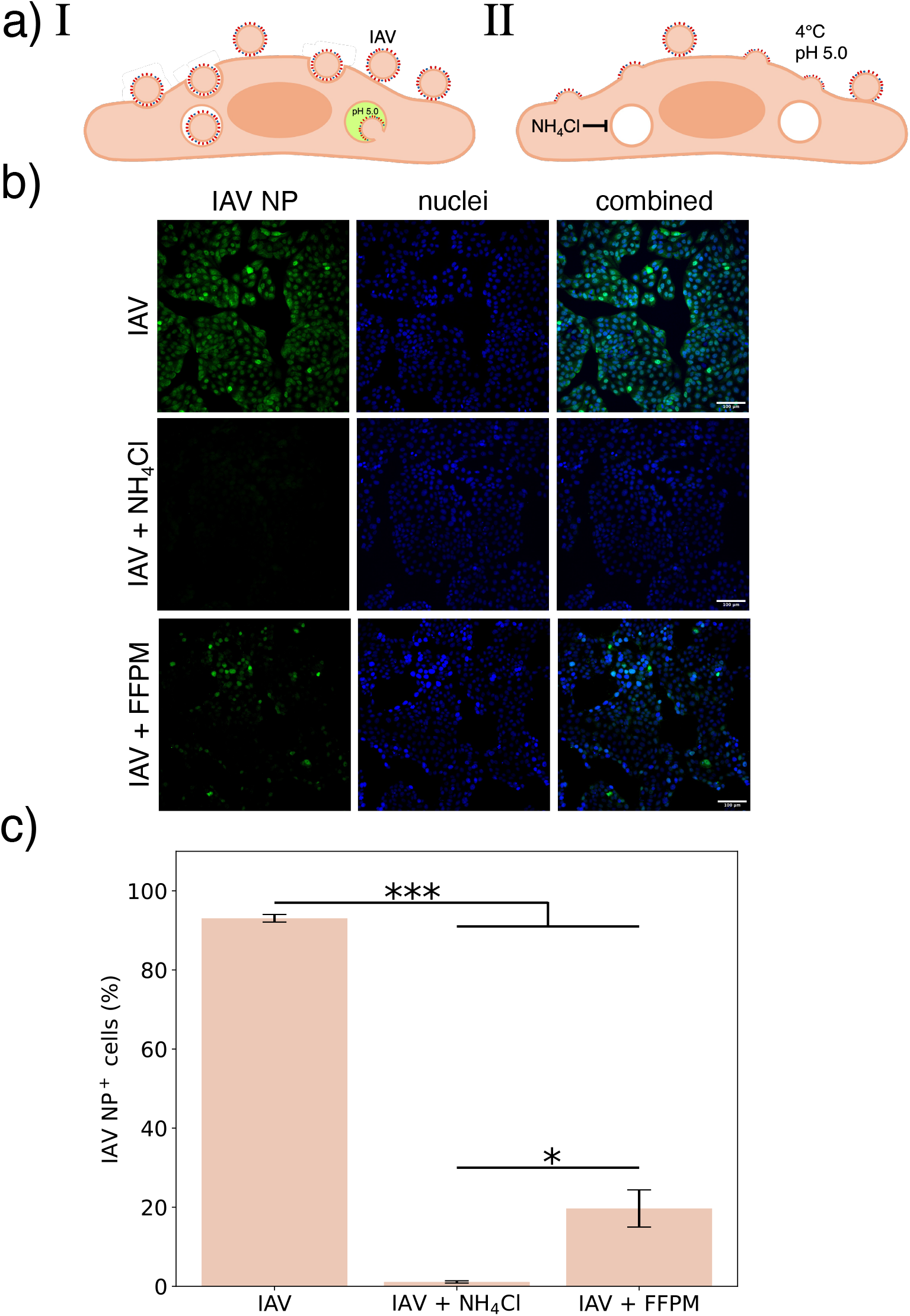
IAV forced fusion at the plasma membrane of cells. **a**) Schematic figure illustrating (**I**) the canonical endocytic pathway for IAV entry and (**II**) forced fusion at the plasma membrane (FFPM). Here, IAV was bound to cells at 4°C to inhibit internalization, followed by a short incubation in a pH 5.0 buffer to trigger fusion. NH_4_Cl was used before and after fusion to prevent fusion triggering within endosomes. Immunostaining of A549 cells for IAV nucleoprotein (IAV NP) 6 hours post-infection was used to assess productive cell infection. **b**) Representative images of cells infected with IAV (top row), in the presence of NH_4_Cl (middle row) and IAV FFPM (bottom row). Fluorescently green cells (left column) indicate cells infected with IAV. **c**) Quantification of IAV NP-positive cells 6 hours post-infection. NP positivity was defined as a nuclear IAV NP intensity above uninfected cells, and the percent NP-positive nuclei was calculated in each sample. A one-way ANOVA was used to determine statistically significant differences between groups. Bar plot shows mean of each group ± standard error (n=3). * and *** indicate p-values < 0.05 and < 0.001, respectively.

### Developing an assay for detecting viral content mixing with PMVs

In order to compare membrane environments in the absence of cell metabolic activity and cytoskeletal effects, we employed cell-derived plasma membrane vesicles (PMVs) to approximate the plasma membrane. PMVs preserve the plasma membrane lipid and protein composition to a large extent, which is difficult to reconstruct in synthetic liposomes (15). However, to what extent lipid asymmetry is conserved after vesiculation is debated (16–18). Because PMVs are crowded with cytoplasmic RNA, we could not utilize our genome exposure assay based on loading target vesicles with a nucleic acid-sensitive dye (19). We therefore chose to utilize the fluorescent dye calcein, loaded inside the PMVs at a quenching concentration, similar to what has previously been performed on liposomes (20).

PMV isolation, calcein incorporation, immobilization in microfluidic flow cells, and binding of virus are schematized in **Figure 2a**. Validation and calibration data for PMV isolation and calcein are shown in **Figure 2b-g**. Briefly, based on a bulk quenching curve (**Fig. 2b**), we chose to resuspend PMVs in 30 mM calcein. This produced quenched fluorescence, as permeabilization with 1% Tween 20 yielded a clean dequenching signal via fluorescence microscopy of immobilized PMVs (**Fig. 2c**). Calcein loading efficiency was estimated at 52 ± 2% (mean ± standard deviation) of PMVs that stained positive with a lipophilic dye. Successful loading of PMVs was facilitated by slow freeze-thaw cycles, which increased the percentage of calcein-loaded PMVs undergoing dequenching by 5-fold (**Fig. 2d**), without altering the size distribution of the PMVs (**Fig. 2e**). This is consistent with previous reports on slow freeze-thawing of giant unilamellar vesicles, where dye incorporation was mainly attributed to pore formation, rather than vesicle fusion/fission (21). In order to test the degree to which plasma membrane asymmetry was preserved, we stained PMVs with FITC-annexin V to measure relative outer leaflet phosphatidylserine concentration. Inducing PMV lipid scrambling with ionomycin increased outer leaflet phosphatidylserine 2-fold (**Fig. 2f**), indicating that PMVs retain some degree of the plasma membrane asymmetry even after freeze-thaw treatment. Finally, electron cryomicrographs of the PMVs show roughly spherical particles with sizes consistent with those measured via nanoparticle tracking analysis (**Fig. 2g**).

**Figure 2.**
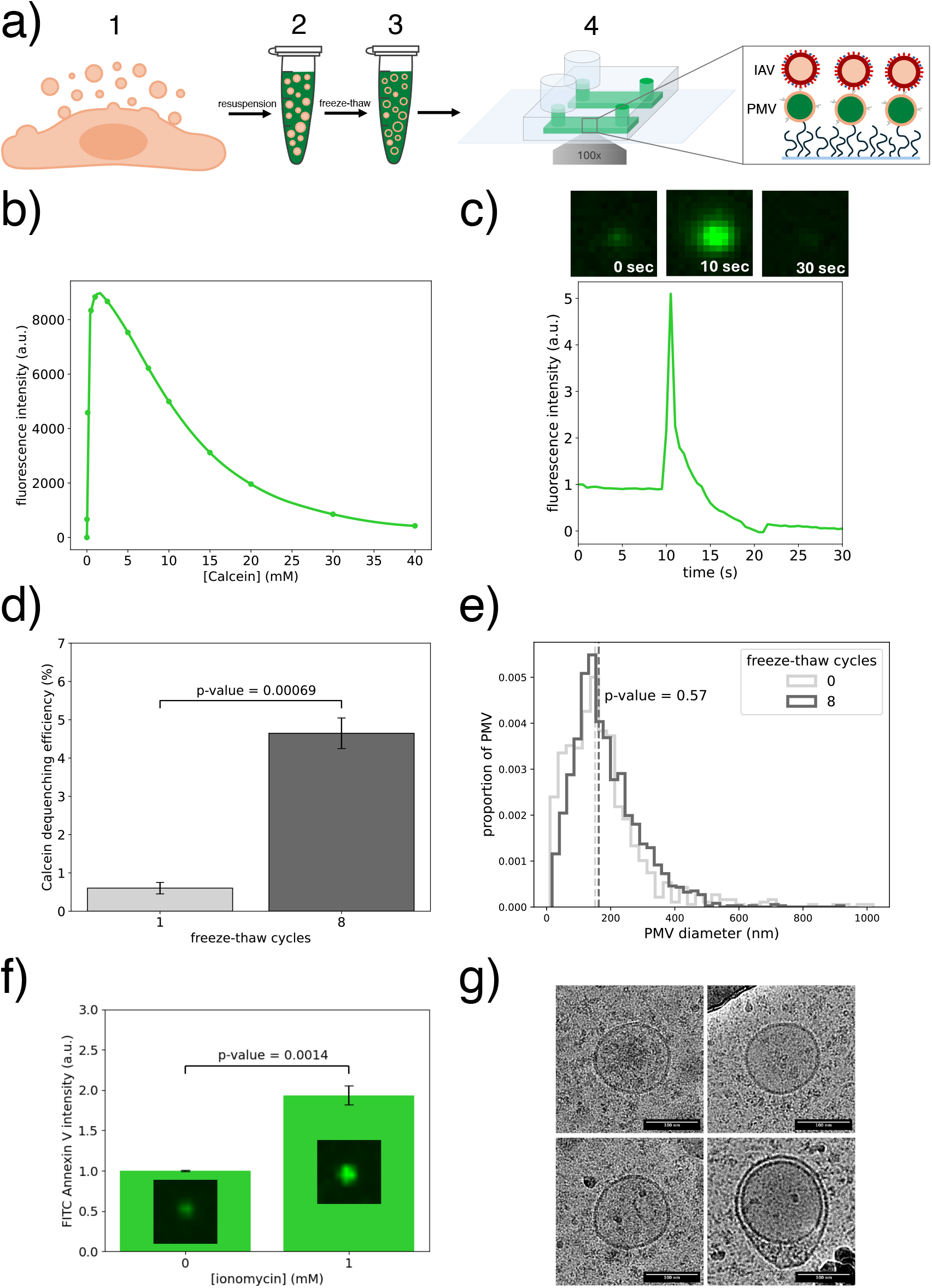
Incorporating quenching concentrations of calcein inside PMVs. **a**) Schematic illustrating the process of PMV isolation, calcein loading, immobilization in a microfluidic flow cell channel, and viral binding: (**1**) Cells were treated with a vesiculation reagent for 2 hours, followed by collection of the PMV-containing supernatant. (**2**) PMVs were pelleted by ultracentrifugation and resuspended in a 30 mM calcein buffer. (**3**) Bulk calcein was incorporated inside the PMVs through 8x slow freeze-thaw cycles. (**4**) Biotinylated PMVs loaded with calcein were immobilized inside neutravidin-coated microfluidic flow cells. Influenza virus was then allowed to bind to PMVs for 5 minutes before unbound virus was washed out. Inset illustrates complete setup prior to triggering fusion. **b**) Bulk calcein quenching curve. Maximal intensity was estimated at 1.6 mM. **c**) Calcein-positive PMV dequenching trace example. The sharp increase followed by complete loss of intensity is interpreted as detergent-facilitated pore formation in the PMV membrane and consequent calcein leakage. Insets above show a single PMV intensity before (0 sec), during (10 sec) and after leakage (30 sec). **d**) Percentage (mean ± standard error) of calcein-positive PMVs that underwent dequenching after detergent poration. A two-sample t-test was conducted to determine a statistically significant difference between number of freeze-thaw cycles for three repeats. **e**) PMV size distribution before and after 8x slow freeze-thaw cycles, determined by nanoparticle tracking analysis. Statistical significance was assessed using a Mann-Whitney U test. **f**) PMVs treated for 5 hours at room temperature with 1 mM ionomycin, followed by labelling with FITC-Annexin V. Data show the FITC intensity (mean ± standard error) of single PMVs from three repeats, and statistical significance was assessed by a two-sample t-test. Inset fluorescence micrographs show sample FITC-Annexin V stained PMVs. **g**) Four sample Cryo-EM micrographs of PMVs after 8x slow freeze-thaw cycles. Note that some PMVs appear multilamellar.

### IAV lipid and content mixing with liposomes versus PMVs

Lipid and content mixing were recorded simultaneously using dequenching of Texas Red-DHPE dye incorporated in the viral membrane and dequenching of calcein dye from the PMV lumen, respectively. 61 ± 4% (mean ± standard error) of Texas Red (TR)-labeled influenza virions were bound to calcein-loaded PMVs (**Fig. 3a**). The rest are likely bound to PMVs that did not successfully incorporate calcein, consistent with the proportion of calcein-negative PMVs after freeze-thawing.

**Figure 3.**
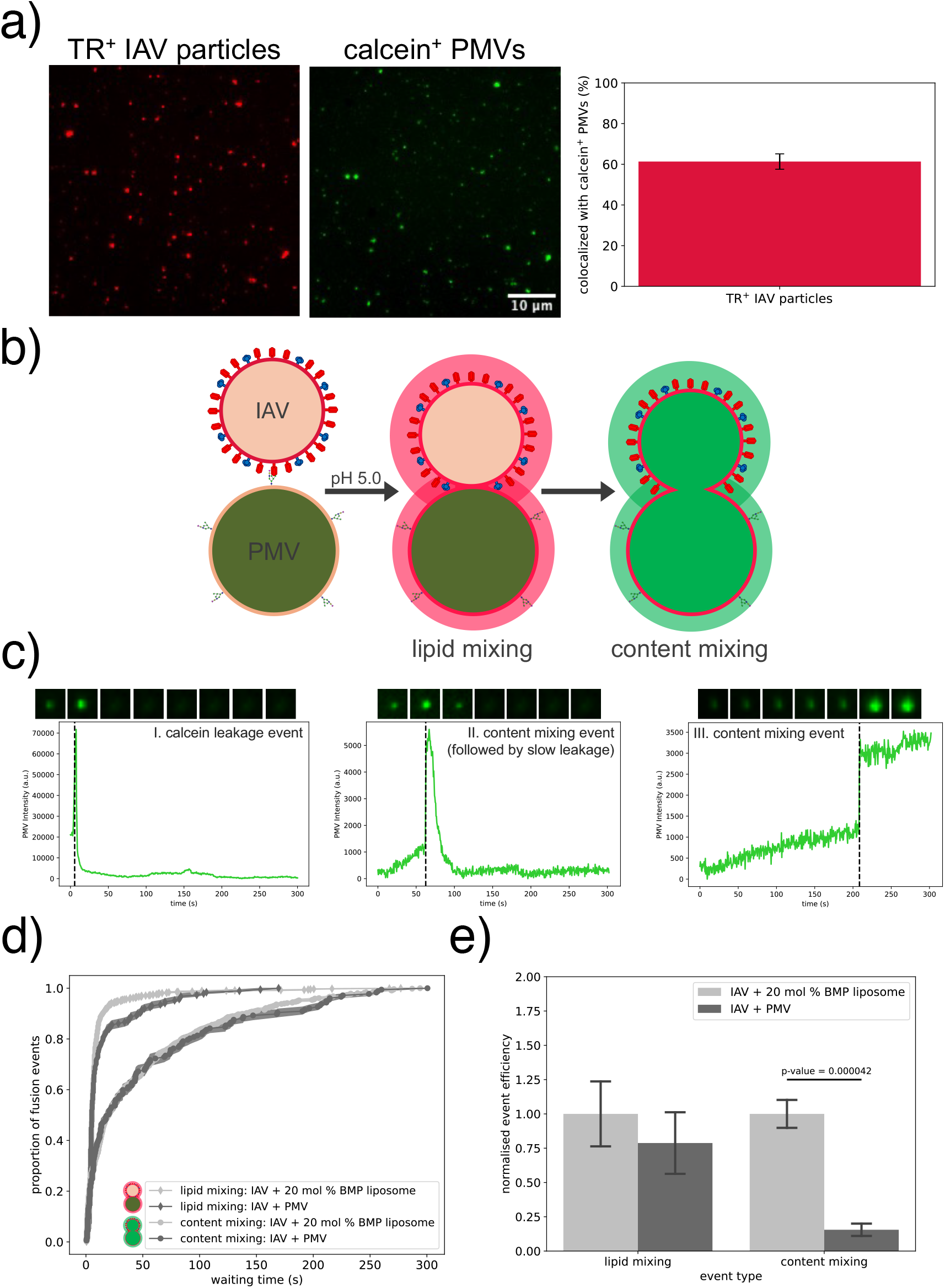
IAV lipid and content mixing with liposomes versus PMVs. **a**) Representative images showing typical channel coverage of Texas Red (TR)-labeled IAV particles (left micrograph) bound to calcein-loaded PMVs (right micrograph). Bar plot shows % (mean ± standard deviation) of TR-positive IAV particles that colocalized with a calcein-positive PMV, from 14 independent channels. **b**) Schematic illustrating a TR-labeled IAV particle bound to a calcein-loaded PMV. Lipid mixing is detected by dilutional dequenching of the TR dye into the PMV membrane. Content mixing is detected by dilutional dequenching of the calcein dye into the viral particle interior. **c**) Calcein intensities over time were plotted for each individual calcein-positive PMV, and those exhibiting a rapid intensity increase after triggering fusion were classified into three groups, based on the length of the intensity peak: “I. calcein leakage event” was characterized by a calcein intensity peak time of <2 seconds, followed by complete loss of calcein signal. “II. content mixing event (followed by slow leakage)” was characterized by a rapid intensity increase, followed by slow intensity decay and “III. content mixing event” was characterized by a rapid intensity increase followed by a stable elevation. Inset images above traces represent PMV intensity profile during the trace. **d**) Lipid and content mixing kinetics at 37°C of IAV bound to 20 mol % BMP liposomes (27 POPC : 20 DOPE : 20 BMP : 30 cholesterol : 2 GD1a : 1 18:1 biotinyl-cap PE) or PMVs, shown as normalized cumulative distribution functions (CDFs) of event waiting times. Filled area around CDFs represent the bootstrapped interquartile range and statistical testing was performed as described in the Methods. **e**) Lipid and content mixing efficiency, calculated as total events recorded / TR-labeled IAV particles detected in the same field of view. Bars show mean ± standard error. A significant difference between content mixing efficiency groups were determined by a two-sample t-test, with p-value annotated on the plot.

The PMV calcein dequenching events obtained after triggering IAV fusion were classified into three types, based on their dequenching profile (**Fig. 3c**). These classes were similar to IAV content mixing observations in liposomes loaded with a quenched fluorescent dye (20). Type I traces were characterized by a rapid increase followed by a rapid drop to background intensity, which we interpret as leakage of calcein into the bulk solution. Using a cytoplasmic diffusion coefficient of 330 µm^2^ per second (22), we estimated the required time for 30 mM calcein to diffuse out of a PMV through a 2.5 nm pore (minimal pore size, chosen as the theoretical size of a calcein molecule) as ∼663 µs. This represents an upper limit on the timescale of calcein depletion from a PMV via free diffusion through a stable pore. Since we observe complete loss of calcein intensity on a >500 ms timescale, the calcein leakage observed as type I traces are unlikely to represent PMV bursting or unrestricted calcein diffusion from a pore/rupture in the membrane. One possibility is that calcein loss occurs through transient membrane pores (23). Nonetheless, since we cannot determine whether a pore was formed between the viral particle and PMV during these calcein leakage events, we excluded these traces from our content mixing analysis. These traces represented ∼5% of all traces detected.

Type II and III traces were characterized by a rapid increase in calcein intensity followed by slow decay or stable elevation, respectively (**Fig. 3c**). We classified these as viral content mixing events, where the intensity peak likely represents calcein dequenching into the viral interior upon pore formation between the viral particle and PMV (20). For type II traces, the slow intensity decay time (>seconds) suggests slow release of calcein into the bulk media. Transient content leakage during HA-mediated membrane fusion has previously been observed and suggested to be related to membrane rearrangements during fusion pore development (24, 25). We observed a similar decay time for similar trace profiles obtained during IAV content mixing with liposomes. Type III traces display a single intensity increase step, indicating an abrupt dequenching event with no subsequent leakage. We cannot differentiate whether this involves full pore opening or a transient pore, but no fluorescence-altering changes to calcein concentration occur after the initial event. The proportion of leaky to non-leaky content mixing events detected overall was approximately 80 : 20 (**Fig. S1**), which is consistent with previous data on cell-cell fusion experiments using HA-expressing cells (24).

Previously we found that the endosomally-enriched phospholipid BMP had a dose-dependent effect on increasing IAV content mixing with liposomes in single-virus fusion assays (3). Liposomes containing 20 mol % BMP yielded indistinguishable content mixing kinetics whether mixing was measured with calcein or the nucleic-acid binding dye DiYO-1 used in our prior work (3; **Fig. S2**). Our previous work has also showed that IAV lipid mixing with liposomes is robust to the target membrane lipid composition (3). Here we find that IAV lipid mixing with PMVs is slightly slower than with synthetic liposomes (**Fig. 3d**), overall similar to our prior results on IAV lipid mixing with endosomal membranes (26). It is possible that protein and/or glycan components of these biological membranes sterically hinder tight apposition after binding, or this slight but noticeable effect could be due to entirely lipidic factors. Moreover, we find that IAV content mixing with PMVs is approximately 6-fold less efficient than with liposomes containing a first-order endosomal mimetic composition (**Fig. 3e**). This difference in fusion efficiency is striking and reproducible, establishing that a large portion of the reduced IAV infection efficiency at the plasma membrane can be assigned to a reduction in fusion efficiency.

### Manipulating the PMV lipid composition and assessing its effect on influenza fusion

Having demonstrated that IAV fusion to the plasma membrane compared to endosome-mimicking liposomes results in slower lipid mixing kinetics and impaired fusion efficiency, we set out to test whether this was caused by the lack of BMP in the plasma membrane. We previously showed that BMP confers a dose-dependent enhancement in IAV content mixing with synthetic liposomes (3), and wanted to test whether supplementing PMVs with BMP would result in a similar enhancement of IAV fusion. First, we attempted to directly supplement the PMVs with exogenous BMP, by estimating the total phospholipid content of the PMV sample (see **supplementary methods item 1**) and directly adding BMP in suspension to the sample. Overall, we observed a 9-fold decrease in content mixing efficiency when supplementing the PMVs with BMP directly, which was constant over a range of 1-20 mol % BMP (**Fig. S3a**). Moreover, supplementing the PMVs with 20 mol % POPC also resulted in the same decrease in content mixing efficiency (**Fig. S3a**). Consequently, we detected too few events to plot any meaningful CDF curves, for estimating content mixing kinetics. Incorporation of phospholipids into preformed vesicles is a complex process and could lead to tighter lipid packing in the outer leaflet (**Fig. S3b**), which has inhibited membrane fusion in other systems (27). Therefore, we opted for the better validated method of pre-loading methyl-α-cyclodextrin (MαCD) with BMP (MαCD-BMP) and delivering BMP to the PMVs using MαCD-assisted lipid exchange. MαCD-based delivery of SM to the outer leaflet of the plasma membrane has been previously described (28), and we have here extended this to BMP. Treating the PMVs with MαCD-BMP resulted in strong anti-BMP immunostaining in the PMVs, supporting effective delivery (**Fig. 4d**). Interestingly, we found that PMVs treated with BMP-MαCD had faster (p < 0.05 with both the KS test and rank sum test, see **Table 1**) lipid mixing kinetics with IAV (**Fig. 4a**). However, incorporating BMP into PMVs did not result in faster content mixing kinetics (**Fig. 4b, Table 1**) or higher content mixing efficiency (**Fig. 4c**), suggesting that BMP alone is not sufficient to explain overall reduced IAV fusion to the plasma membrane (**Fig. 3e**).

**Table 1.**
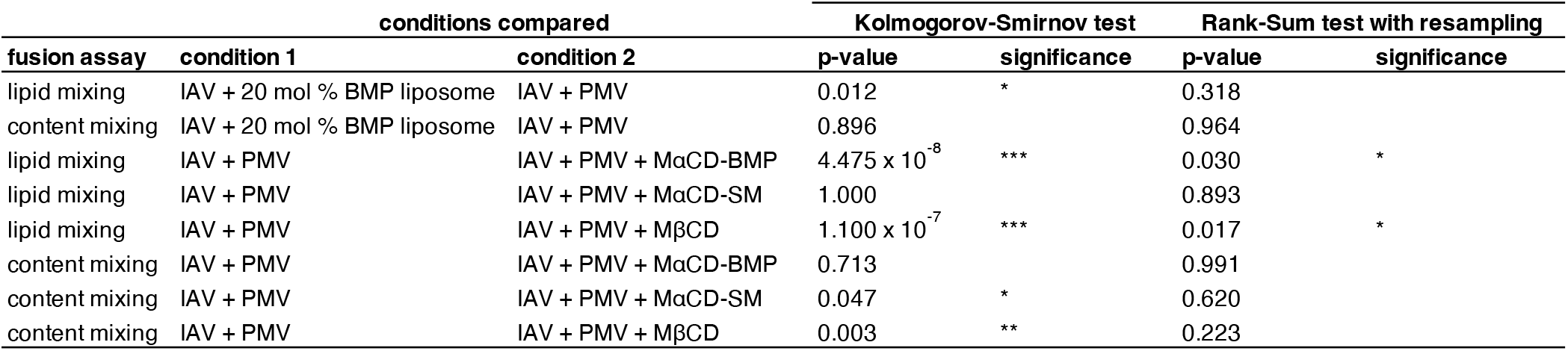
Significance testing for fusion kinetics between IAV and different target vesicles. Significance testing was performed between single-event waiting time distributions for each condition, corresponding to the CDFs in **Fig. 3d, Fig. 4a** and **Fig. 4b**. Kolmogorov-Smirnov (KS) tests with Bonferroni multiple hypothesis corrections were performed between the aggregate waiting times for each condition. To better capture variation between flow-cell channels, bootstrapped Wilcoxon Rank Sum Tests were performed with 10 000 bootstrap resamples across flow cells, and Bonferroni-corrected p-values are reported again. Annotations *, ** and *** signify p-values below 0.05, 0.01 and 0.001, respectively.

**Figure 4.**
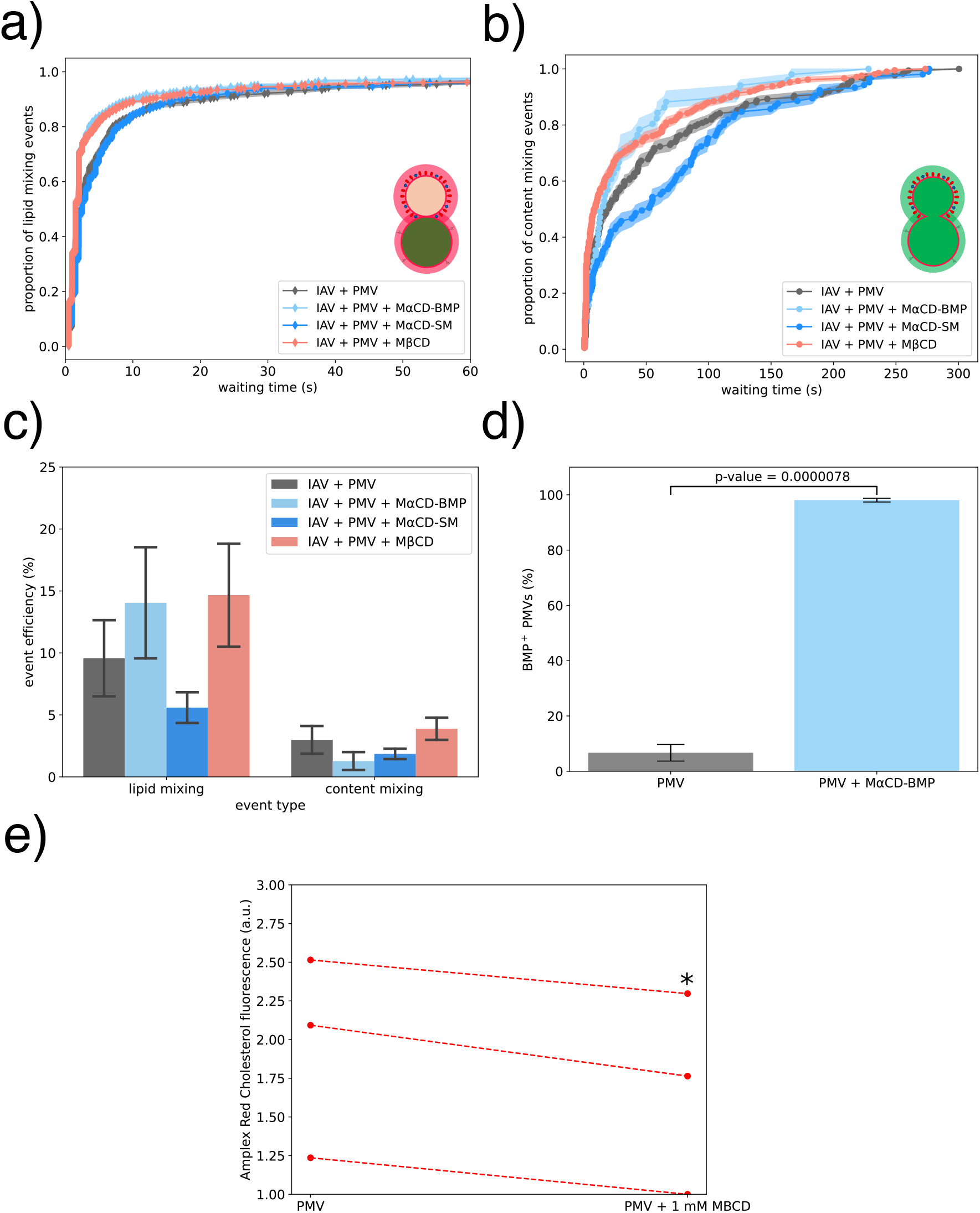
Effect of membrane perturbations on IAV lipid and content mixing with PMVs. **a**) Lipid mixing kinetics between IAV and PMV that have been treated to modify membrane composition as indicated. Filled area around CDFs represent the bootstrapped interquartile range, and statistical significance was performed as described in the Methods. Only the first 60 seconds of event waiting times are shown to better highlight differences between CDFs, but calculations are performed on the full 300 seconds. **b**) Content mixing kinetics of IAV with PMVs that have been treated to modify their membrane composition. The full 300 second time course is shown. **c**) Lipid and content mixing efficiency, calculated as total events recorded / TR-labeled IAV particles detected in the same field of view. One-way ANOVAs calculated separately for the lipid (p-value = 0.38) and content mixing (p-value = 0.42) efficiency groups revealed no significant difference between groups. **d**) Delivery of BMP to PMVs using MαCD quantitated via immunolabeling. Bar plot shows the % (mean ± standard error) of BMP-positive spots that colocalized with PMV-positive spots measured on three separate channels, with the p-value from a two-sample t-test annotated on the plot. **e**) Cholesterol content of PMVs treated with MßCD, as determined by a cholesterol esterase assay. The baseline measured value was variable between repeats, but the 14 ± 3% (mean ± standard error) decrease in cholesterol was statistically significant (p<0.05) via a paired two-sample t-test.

We previously showed that phosphatidylserine (PS) has a similar effect to BMP in enhancing content mixing efficiency in liposomes (3). Native plasma membrane has high inner leaflet, but low outer leaflet PS content (29), so we hypothesized that loss of lipid asymmetry might increase IAV content mixing efficiency in a similar fashion. Treating PMVs with ionomycin did result in increased outer leaflet PS exposure (**Fig. 2f**). However, it did not affect content mixing kinetics (**Fig. S4a**) or efficiency (**Fig. S4b**).

The outer leaflet of the plasma membrane is characterized by a high content of cholesterol, SM and fully saturated phospholipids, resulting in lower membrane fluidity, tighter lipid packing, and a liquid-ordered-like character (30). Conversely, the inner leaflet has a much higher degree of unsaturation. We hypothesized that the liquid-ordered-like composition of the plasma membrane outer leaflet may slow or restrict viral membrane fusion. Interestingly, the asymmetry itself appears not to be a major restrictive factor, as treatment of PMVs with ionomycin constitutes a good control for overall asymmetry and not just PS exposure: ionomycin-induced scrambling did not significantly change content mixing kinetics (**Fig. S4a**) or efficiency (**Fig. S4b**). We found that reducing the cholesterol content of PMVs via MßCD extraction resulted in faster IAV lipid and content mixing kinetics (**Figs. 4a, 4b, and Table 1**), without affecting efficiency (**Fig. 4c**). Although the enhancement of content mixing kinetics between IAV and MßCD-treated PMVs did not reach statistical significance, it was notable and expected given the rate enhancement of the upstream lipid mixing process between these two partners. This relatively mild MßCD treatment reduced PMV cholesterol content by 14%, as measured by a cholesterol oxidase activity (**Fig. 4e**), consistent with previous reports on whole cells (31).

Cholesterol has been shown to have a strong effect in promoting influenza infectivity and full fusion both within the viral envelope and in synthetic target liposomes. Such an effect may occur potentially by modulating the free energy of highly curved fusion intermediates (20, 25, 32–34). Interestingly, endosomal membranes that constitute the physiological site of IAV fusion have lower cholesterol content than the plasma membrane (9). We believe that the enhancement in fusion achieved by extracting cholesterol from PMVs is related to plasma membrane organization and the liquid-ordered-like nature of the outer leaflet rather than the direct physical properties of cholesterol at the fusion interface, which are likely fusion-promoting. Interestingly, prior results in synthetic liposomes have shown that cholesterol content affects the proportion of content mixing events that are non-leaky (20, 35). We found that reduction in PMV cholesterol content did not affect the proportion of non-leaky content mixing events (**Fig. S1**), supporting the notion that a modest reduction in PMV cholesterol content is not acting by modulating the free energy barriers for transition between curved fusion intermediates. We recognize that cholesterol depletion from the outer leaflet can cause more complex changes in leaflet structure and composition. However, since fusion was unaffected by ionomycin-induced scrambling (**Fig. S4**), the MßCD effect on fusion cannot be attributed to intra-membrane redistribution alone.

Since liquid-ordered phase components appear to affect influenza fusion, we also tested perturbations to the liquid-ordered plasma membrane component sphingomyelin (SM). Delivering additional SM to the outer leaflet of PMVs using MαCD pre-loaded with SM (MαCD-SM) did not alter lipid mixing kinetics (**Fig. 4a**) and affected content mixing kinetics in a fashion that was large in magnitude (**Fig. 4b**) but highly heterogeneous across biological repeats and ultimately not statistically significant (p > 0.5 via rank sum test bootstrapped across flow cells, **Table 1**). Content mixing efficiency was not significantly altered (**Fig. 4c**). These results suggest there is not a strong dose-dependent effect of SM concentration on fusion and thus that the lipid is not substantially affecting the free energy of curved fusion intermediates. Because SM is normally present at relatively high mole fractions in the plasma membrane outer leaflet, it may be that large-scale changes affect full fusion kinetics and that such effects are responsible for the heterogeneity seen in the MαCD-SM treated samples, but this remains speculation.

### PMV membrane perturbations affect membrane order or fluidity

Given the strong effect of perturbations to liquid-ordered membrane components on IAV fusion (**Fig. 4a and b**) and prior work showing that host restriction factors such as IFITM3 and Serinc5 alter membrane order and stiffness (36, 37), we hypothesized that similar effects may control influenza fusion with the plasma membrane. We assessed this by measuring the Laurdan fluorescence spectra in liposome and PMV preparations pre-labeled with C-Laurdan under the conditions where we also measured fusion. Interestingly, Laurdan in the membrane of the PMVs exhibited a substantially blue-shifted fluorescence spectrum compared to the 20 mol % BMP liposomes (**Fig. 5a**). This is suggestive of greater membrane order or lower fluidity (38) and may explain some of the difference in IAV fusion kinetics and efficiency between the 20 mol % BMP liposomes and PMVs. To quantitate this, we calculated the Laurdan generalized polarization (GP) of the 20 mol % BMP liposomes and PMV samples treated with different soluble factors from their respective Laurdan emission spectra (39).

**Figure 5.**
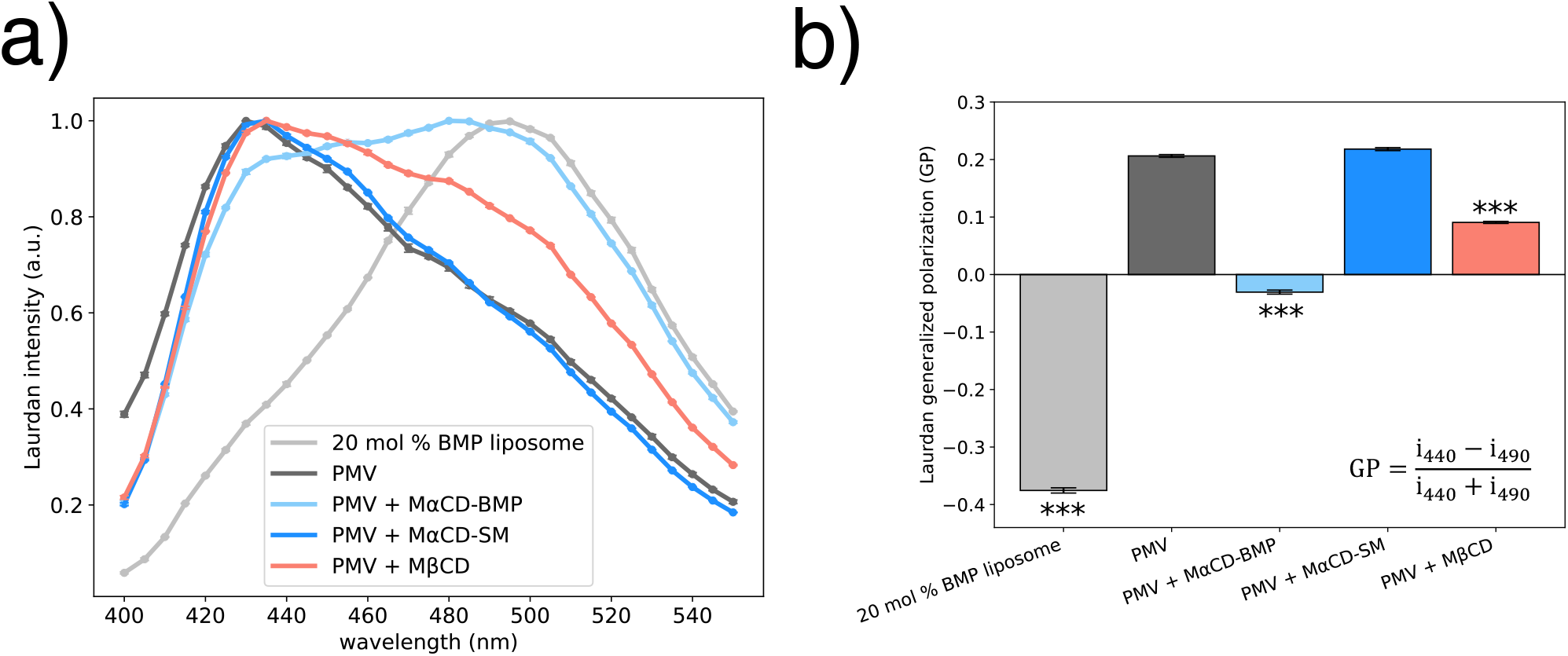
Membrane order of liposomes and PMVs. **a**) Laurdan fluorescence emission spectra at 37°C of 20 mol % BMP liposomes and PMVs with indicated membrane modifications corresponding to those tested in **Fig. 4**. Fluorescence intensity is normalized to 1 for the peak intensity and each point represents mean ± standard error from three repeats. **b**) Laurdan generalized polarization (GP) of 20 mol % BMP liposomes and PMVs, calculated from the fluorescence emission spectra given in a). The equation for calculating GP values is displayed, where i_x_ signifies intensity at wavelength x (39). Vesicles with a significantly different GP value from the PMV sample, determined via a one-way ANOVA and Tukey HSD post-hoc test, are labeled with ***, denoting p < 0.001.

Compared to PMVs, cholesterol-depleted PMVs exhibited a significantly lower Laurdan GP value (**Fig. 5b**), suggesting that a modest reduction in cholesterol content results in overall higher membrane fluidity in PMVs (39). This is consistent with the well-known role of cholesterol in regulating plasma membrane order and fluidity (40). Cholesterol’s association with SM or other saturated lipids is thought to be a key factor in liquid-ordered phase formation and the observed ordering of plasma membrane outer leaflets (41–43). It is thus not surprising that depleting cholesterol from the high-cholesterol, high-SM PMVs would reduce membrane order. This is associated with enhanced fusion. Such an effect is distinct from the well-documented effect whereby cholesterol promotes fusion in non-phase-separating liposomes (32). However, increasing PMV SM content with MαCD-SM did not affect membrane order (**Fig. 5b**), suggesting that increasing the plasma membrane SM content beyond physiological levels have limited effects on the overall plasma membrane order and fluidity. Interestingly, both BMP incorporation and cholesterol depletion in PMVs resulted in faster lipid mixing kinetics (**Fig. 4a**) and higher membrane fluidity (**Fig. 5b**), suggesting a potential influence of target membrane fluidity on stalk formation. This is separate from the previously observed effect of BMP on IAV full fusion in simple liposomes (3).

### IAV forced fusion at the plasma membrane of cholesterol-depleted cells

Since treating PMVs with MßCD increased IAV fusion kinetics (**Fig. 4a and b**), we hypothesized that treatment of a lung epithelial cell line with MßCD prior to infection would increase IAV productive infection via FFPM. Representative images are rendered in **Fig. 6a**, and IAV NP immunopositivity of all imaged cells is plotted in **Fig. 6b**. Pre-treatment of the A549 cells with 1 mM MßCD resulted in a 2.4-fold increase in infected cells after FFPM (**Fig. 6b**). This suggests that cholesterol extraction from the plasma membrane reorganizes the membrane in a way that promotes IAV fusion at the plasma membrane and consequently productive cell infection. We speculate that the observed increase in productive infection after a modest cholesterol extraction is related to decreased plasma membrane order and/or fluidity and disruption of liquid-ordered domains (43). Increasing the MßCD dose to 5 mM resulted in loss of cell-cell adherence in the cell monolayer (**Fig. S5a**). This higher concentration reduced the number of infected cells both when cells were infected through the canonical endosomal route and FFPM (**Fig. S5b**). We attribute this to impaired IAV binding to cells (44) and reduced cell viability (45), as previously shown, rather than inhibition of infection.

**Figure 6.**
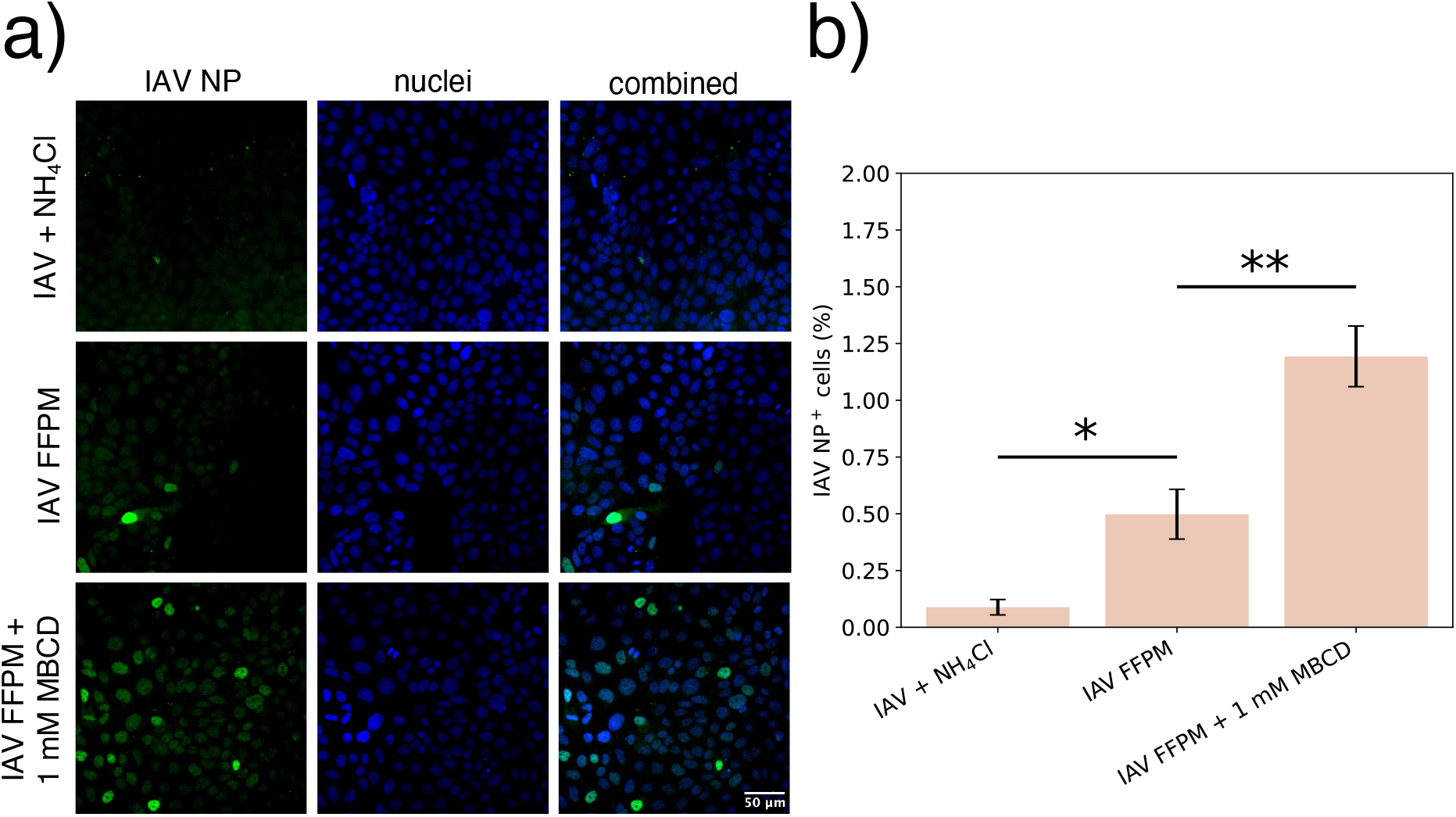
IAV FFPM with A549 cells after MßCD treatment. **a**) Representative images of cells infected with IAV in the presence of NH_4_Cl (top row), IAV FFPM (middle row) and IAV FFPM after treating cells with 1 mM MßCD for 30 min at 37°C (bottom row). Fluorescently green cells (left column) indicate cells infected with IAV. **b**) Quantification of IAV NP-positive cells 5 hours post-infection. NP positivity was defined as a mean nuclear NP intensity above uninfected cells, and the percent NP-positive nuclei was calculated in each sample. A one-way ANOVA and Tukey HSD post-hoc test was used to determine statistically significant differences between groups, with significantly different groups labeled on the plot. Bar plot shows mean of each group ± standard error from three repeats. * and ** indicate p-values < 0.05 and < 0.01, respectively.

## Discussion

Influenza virus has long been known to be capable of forced fusion at the plasma membrane, albeit at lower infection efficiency than physiological fusion within endosomes. Here we demonstrate that a portion of this lower infection efficiency is due to membrane differences: respiratory epithelial cell plasma membranes are less permissive to influenza fusion than are endosome-mimetic liposomes. To our surprise, incorporating the endosome-enriched phospholipid BMP into the plasma membrane was not sufficient to promote influenza full fusion, underscoring the complexity of biological cell membranes versus simple synthetic liposomes (3). We hypothesize that this may be related to the liquid-ordered-like nature of the plasma membrane outer leaflet. Either extraction of cholesterol from plasma membrane vesicles or delivery of BMP speeds lipid mixing and introduces a concomitant less-ordered component to the Laurdan fluorescence spectrum. Under these conditions we did not, however, observe a significant change in full fusion rates or efficiencies. It is possible that such changes would be observed with greater changes in the mol % of BMP or cholesterol, especially given the ability of MßCD extraction of cholesterol to promote influenza infection at the plasma membrane.

Alternatively, the observed effects could be due to a process that affects only the first stages of tight apposition and lipid exchange in membrane fusion. Lateral inhomogeneity in the plasma membrane could cause this if some regions of the plasma membrane were impermissive to fusion and others were permissive, or it could be due to changes in membrane deformation free energies at the site of fusion. The findings presented here do not definitively assign a mechanistic basis, but they establish that differences in influenza infection efficiency across subcellular compartments can be largely assigned to membrane factors and that some of these factors are related to the liquid-ordered nature of the plasma membrane outer leaflet.

There exist abundant data on how target membrane composition can affect enveloped viral entry, but mechanistic studies of this nature primarily focus on either synthetic membranes or positive modulators of fusion (20, 32, 46–51). Here, we show that the plasma membrane lipid composition can restrict entry of an endosome-fusing virus. In combination with prior work on endosomal membrane factors that promote fusion (3, 5–7) and that HIV-1, but not influenza, preferentially fuses to membrane domain phase boundaries (10), these results raise the question of whether the plasma membrane is broadly less permissive to viral fusion than endosomes, or whether endosome-fusing viruses have adapted to their subcellular niche.

## Methods

### Cell culture and virus strain

A549 cells were maintained in complete DMEM (cDMEM): DMEM, 4.5 g/L glucose with Glutamax (Gibco) supplemented with 10% (v/v) Fetal Bovine Serum (Gibco) and 100 units/ml Penicillin and 100 µg/ml Streptomycin (Gibco), at 37°C with 5% CO_2_. Cells were kept below 25 cell passages. The X-31 A/Aichi/68 (H3N2) IAV strain was purchased from

AVS Bio as sucrose-purified allantoic fluid and stored at -80°C, until use. To measure lipid mixing, the IAV membrane was labeled with Texas Red-DHPE (Invitrogen), as described elsewhere (19).

### Fluorescence microscopy

All fluorescence imaging was carried out on a Zeiss Axio Observer epifluorescence microscope using a Lumencor Spectra-X LED light source, a 100x, 1.4 NA objective, and two Andor Zyla sCMOS cameras with a dichroic beam splitter to permit simultaneous imaging in two channels. Micromanager 2.0 was used for image acquisition.

### IAV forced fusion at the plasma membrane

A549 cells were seeded at 2 x 10^5^ per well in an 8-well cell culture chamber (Sarstedt). The following day, the cells were ∼60% confluent and preincubated for 10 minutes at 37°C in cDMEM containing 30 mM NH_4_Cl before cooling to 4°C on ice for 10 min. Cells were then incubated in cooled cDMEM containing IAV (with an estimated multiplicity of infection of 2.0) and 30 mM NH_4_Cl on ice for 20 min with gentle agitation to allow virus adsorption. Media was then exchanged with pH 5.0 reaction buffer (10 mM Sodium Phosphate, 90 mM Sodium Citrate, 150 mM NaCl, adjusted to pH 7.4 or 5.0 as stated) pre-warmed to 37°C and incubated on a 37°C heat block for 1 minute, to trigger IAV fusion at the plasma membrane and recommence endocytosis. Wells were subsequently washed once and incubated in cDMEM containing 30 mM NH_4_Cl. One hour before fixing, infected cells were incubated in cDMEM containing 30 mM NH_4_Cl and 1:1500 SYTO Deep Red Nucleic Acid stain (Invitrogen). 6 hours post-infection, cells were fixed in a 4% (w/v) paraformaldehyde solution for 15 minutes, followed by permeabilization with a 0.2% (v/v) Triton X-100 solution for 15 minutes. Non-specific antibody binding was blocked by 1% (w/v) BSA for 1 hour, before cells were incubated with a rabbit polyclonal anti-influenza Nucleoprotein antibody (Invitrogen) diluted 1:1000 in 0.1% (w/v) BSA solution overnight at 4°C. The following day, the cells were incubated for 1 hour in the dark with a chicken anti-rabbit secondary antibody conjugated to Alexa Fluor 594 diluted 1:1000 in 0.1% (w/v) BSA. Subsequently, cells were dried and mounted using ProLong mounting media (Invitrogen). The cells were washed 2x with PBS between each step following 6 hours post-infection, and carried out at room temperature, unless otherwise stated. Cells were imaged with a 0.8 NA 20x Zeiss lens, with overlayed images of the nuclear channel (Chroma Far Red filter set) and IAV nucleoprotein (IAV NP) channel (TR filter set). The nuclear IAV NP intensities for each cell was analyzed in Icy Bioimager (52), using an image analysis pipeline (available at https://github.com/kassonlab/Icy_Bioimager_IAV_NP_Nuclear_Intensity_analysis). In short, the nucleus of each cell was segmented and the average IAV NP intensity per nucleus was measured. In Python, an intensity threshold for classifying a cell as IAV NP-positive (IAV infected) was estimated as the intensity of which 99.9% of mock infected cell nuclei fell below. Then the number of IAV NP-positive nuclei (nuclei with a NP intensity above the threshold) over total nuclei was calculated to estimate the % of infected cells. Five images were taken of each well, and each biological repeat (n) consisted of one individual well. All cells per image were analyzed and in total >38,000 cells were analyzed for nuclear IAV NP intensity.

### PMV formation and purification

3x ∼90% confluent T175 flasks seeded with A549 cells were incubated with GPMV buffer (150 mM NaCl, 2 mM CaCl_2_, 20 mM Hepes, adjusted to pH 7.4) supplemented with 25 mM PFA and 2 mM DTT for 2 hours at 37°C, followed by tapping the flasks to aid PMV detachment. The PMV suspension was centrifuged at 150xg for 5 min to remove cell debris and pellet giant PMVs. The supernatant containing smaller PMVs was collected and concentrated by ultracentrifugation at 38 000xg for 1 hour, followed by resuspension of the PMV pellet in calcein buffer (10 mM Hepes, 150 mM NaCl, 30 mM calcein, adjusted to pH 7.4). Next, 50 µM PE-Biotin was added to the PMV sample and incubated shaking at 800 rpm for 30 min, to promote lipid incorporation. PMVs were then subjected to 8x cycles of slow freezing at -80°C, followed by slow thawing at room temperature, as suggested elsewhere for efficient dye exchange in giant unilamellar vesicles (21). No liquid nitrogen or water was used for freeze-thawing the PMVs. PMV size quantification was performed via nanoparticle tracking analysis using a Nanosight instrument (Malvern Panalytical) with a 1:2000 dilution of samples in pH 7.4 reaction buffer.

### Liposome preparation

Liposomes were prepared as described elsewhere (19). In short, 2 mM lipids were mixed and dried into a lipid film by nitrogen effusion, followed by resuspension in pH 7.4 reaction buffer. After vigorous vortexing of the lipid mixture, the liposomes were subjected to 5x freeze-thaw cycles in liquid nitrogen. Unilamellar liposomes with a nominal diameter of 200 nm were further generated through extrusion. Freshly extruded liposomes were diluted in calcein buffer to reach 30 mM bulk concentration, followed by 8x slow freeze-thaw cycles, as described above for PMVs. The 20 mol % BMP liposomes consisted of 27 mol % POPC, 20 mol % DOPE, 20 mol % bis(monoacylglycerol)phosphate (BMP), 30 mol % Cholesterol, 2 mol % GD1a and 1 mol % 18:1 Biotinyl Cap PE. All lipids were purchased from Avanti Polar Lipids.

### Microfluidic flow cell preparation and PMV treatment with membrane-modifying factors

Microfluidic flow cells were prepared as described previously (53). In short, a PDMS cast containing channels was attached to a glass coverslip, and the glass surface was coated with PLL-PEG-Biotin, followed by neutravidin. Biotinylated PMVs or liposomes were added to the flow cell channel and allowed to bind to the neutravidin-coated surface overnight at 4°C. The next day, unbound vesicles and bulk calcein was washed out. If the PMV membranes were designated to be modified, 10 µl of the membrane-modifying factor was added to the flow cell channel, followed by incubation at 37°C for the indicated time. The factor was then washed out of the channel. Lastly, 5 µl Texas Red (TR)- labeled IAV suspension was added to the wells, incubated for 5 minutes at room temperature, followed by washing out unbound virus. All washes were done with pH 7.4 reaction buffer.

### Colocalization analysis to validate BMP delivery

PMVs treated with MαCD-BMP were pre-labeled with the lipophilic dye CellMask Deep Red Plasma Membrane Stain (Invitrogen) at 2x the recommended concentration for cell labelling, followed by shaking the sample at 800 rpm for 30 min to promote dye incorporation. After MαCD-BMP treatment in the flow cells, PMVs were labeled against BMP by incubating them with a mouse anti-BMP antibody diluted 1:1000 (Sigma-Aldrich) for 60 min, followed by washing out the antibody and incubating with a FITC-conjugated anti-mouse secondary antibody (Abcam) for 10 min, before washing unbound antibody. All steps were done at room temperature and antibodies were diluted in pH 7.4 reaction buffer containing 0.1% (w/v) BSA. Labeled vesicles/virus particles imaged in both channels were overlayed in Fiji (54). Detection, colocalization, and quantification of fluorescent spots were performed using the ComDet plugin for ImageJ (https://github.com/UU-cellbiology/ComDet), with a max pixel distance set to 5, spot pixel size to 2 and spot intensity threshold above background to 3 standard deviations.

### Single-virus microscopy in microfluidic flow cells

IAV was bound to immobilized calcein-loaded vesicles inside the flow cell chamber. Rapid buffer exchange to pH 5.0 reaction buffer inside the channel triggered IAV fusion with the vesicles (53). All experiments were performed at 37 ºC. Fluorescence images were acquired at a rate of two images per second, with an exposure time of 200 ms, nominal LED light source power of 4%, and 4×4 pixel binning. Texas Red and calcein intensities were imaged simultaneously using a dichroic beamspliter (Chroma T560lpxr-UF2) and paired cameras. Single-particle TR or calcein intensities were extracted from each image using the ImageJ plugin Spot Intensity Analysis (Nico Stuurman and Ankur Jain, 2018), with spot radius set to 1 and noise tolerance set to 100. Time 0 (when fusion was triggered) was manually set as the frame where the calcein background intensity dropped, corresponding to a local pH change to 5.0 (typically ∼2.5 seconds after commencing buffer exchange). The extracted intensity traces were analyzed for a sharp increase in TR or calcein intensity over time, interpreted as a potential viral lipid or content mixing event, respectively. The Python code for single-particle event analysis is available from https://github.com/kassonlab/single_particle_event_analysis/. Lastly, the potential lipid or content mixing events were manually curated for false positive events, before the waiting time from fusion trigger (pH 5.0 buffer exposure) to the event (sharp increase in fluorescence intensity) for all events detected were plotted as a cumulative distribution function (CDF).

Statistical testing for differences in lipid mixing or content mixing kinetics was performed as follows. We evaluated two error models, one in which virus-to-virus differences were the major source of variation and one in which flow-cell-to-flow-cell differences were the major source of variation. Differences in fusion kinetics were evaluated via a two-sample Kolmogorov-Smirnov (KS) test in the first instance. In the second instance, data from each microfluidic flow cell were kept separate and bootstrap resampling was performed across flow cells. Differences between complied CDFs from each bootstrap sample were evaluated using a rank sum test, and the overall p-value was calculated as the fraction of bootstrap resamples that were found significantly different. Compared to our prior work on synthetic liposomes, we observed a greater degree of heterogeneity between flow cell channels. This may be due to a lower density of PMV:influenza virus conjugates captured per field of view, or it may stem from biological heterogeneity in the plasma membrane vesicle samples themselves. Because differences between channels accounted for more variation in fusion wait times than differences between individual particles in this dataset, we used bootstrapped rank sum tests across channels as the definitive measure of significance. Both significance measures are reported for each pair of conditions compared in **Table 1**.

### C-Laurdan imaging

Vesicles were labeled by adding 10 µM C-Laurdan to vesicle sample, followed by 30 min incubation at room temperature with 800 rpm shaking. Next, vesicles were diluted 1:5 in Hepes buffer (20 mM Hepes, 150 mM NaCl, adjusted to pH 7.4) and added to wells in a glass bottomed 96-well plate. The bulk vesicle C-Laurdan fluorescence spectra was measured in a Synergy H1 microplate reader with the accompanying Gen5 software, with an excitation wavelength of 348 nm and gain set to 70.

## Data availability

Single-virus fusion data are available from Zenodo at doi:10.5281/zenodo.15085255.

## Acknowledgements

This work was supported by grants from the Knut and Alice Wallenberg Foundation (KAW 2020.0209) and the Swedish Research Council (2023-04674) to P.M.K. Electron cryomicrographs of PMVs were acquired using the Cryo-EM Uppsala facility.

