## Supplemental Methods for "Influenza virus fusion at the plasma membrane is restricted by liquid-ordered membrane components"

To estimate the number of phospholipids in a single PMV, we first calculated the radii of the outer and inner leaflets of the vesicle membrane. The outer leaflet radius ($r_{outer\ leaflet}$) was taken as the mean vesicle radius, as given by the Nanosight nanoparticle tracker of the PMV sample. The inner leaflet radius ($r_{inner\ leaflet}$) was taken as $r_{outer}$ minus half the assumed bilayer thickness (7 nm). Next, the area of the bilayer occupied by proteins was estimated to 25 %, hence the total lipid Bilayer area was adjusted accordingly (Dupuy and Engelman 2008):

$$\text{PMV bilayer area} = 4\pi(r_{\text{outer leaflet}})^2 + 4\pi(r_{\text{outer leaflet}} - 7\ \text{nm})^2 \times (1 - \text{protein content})$$

Next, the total number of phospholipids ($N_{PMV\ phospholipids}$) per PMV was estimated by dividing the PMV bilayer area with the assumed phospholipid area ($A_{phospholipid}$) of 0.513 $nm^2$, as reported elsewhere (Alberts et al. 2002):

$$N_{\text{PMV phospholipids}} = \frac{\text{PMV bilayer area}}{A_{\text{phospholipid}}}$$

To convert from estimated phospholipids per PMV to total phospholipids in the sample ($N_{sample\ phospholipids}$), the $N_{PMV\ phospholipids}$ was multiplied by the PMV concentration ($C_{PMV\ sample}$), as determined by the Nanosight nanoparticle tracker, and the total sample volume ($V_{PMV\ sample}$):

$$N_{\text{sample phospholipids}} = N_{\text{PMV phospholipids}} \times C_{\text{PMV sample}} \times V_{\text{PMV sample}}$$

Lastly, the amount of phospholipids to supplement the PMV to reach a desired mol % ratio of the total phospholipid content was estimated by:

$$N_{\text{phospholipid supplement}} = \frac{N_{\text{PMV phospholipids}} \times \text{ratio}_{\%\ \text{mol}}}{(1 - \text{ratio}_{\%\ \text{mol}})}$$

The volume of a certain phospholipid to add to the PMV sample to reach the desired mol % ratio was calculated by the molar weight of the phospholipid and molarity of the stock solution.

Alberts, Bruce, Alexander Johnson, Julian Lewis, Martin Raff, Keith Roberts, and Peter Walter. 2002. ‘The Lipid Bilayer’. In *Molecular Biology of the Cell. 4th Edition*. Garland Science. https://www.ncbi.nlm.nih.gov/books/NBK26871/.

Dupuy, Allison D., and Donald M. Engelman. 2008. ‘Protein Area Occupancy at the Center of the Red Blood Cell Membrane’. *Proceedings of the National Academy of Sciences of the United States of America* 105 (8): 2848–52. https://doi.org/10.1073/pnas.0712379105.
