## Supplemental Data for "Influenza virus fusion at the plasma membrane is restricted by liquid-ordered membrane components"

**Influenza viral infection at the plasma membrane is restricted by lipid composition**

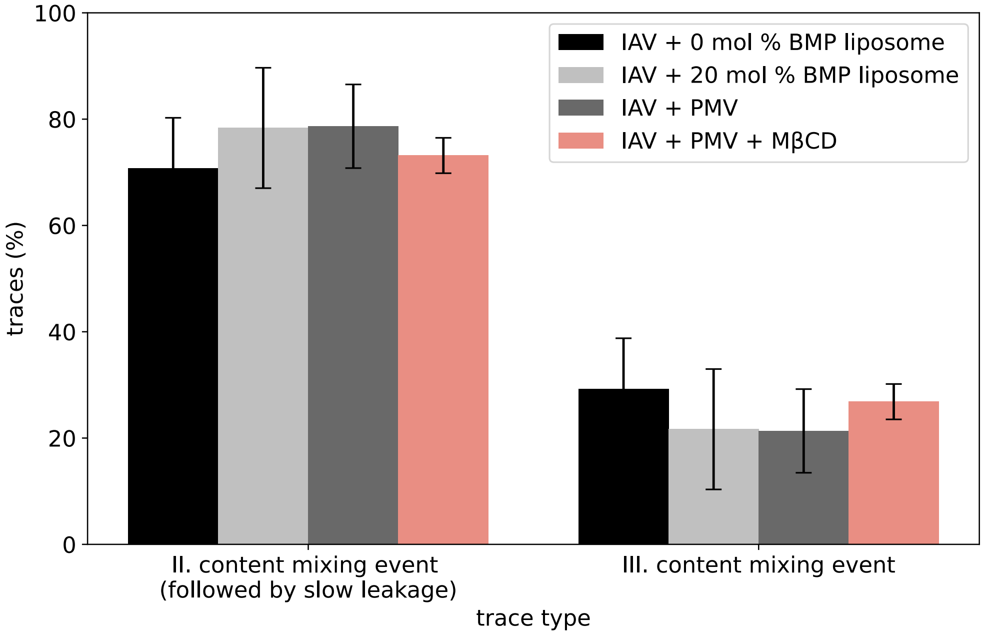


**Figure S1. Distribution of IAV content mixing events classified as Type II or III.** The % of traces which displayed the characteristic calcein intensity followed by a slow decay (Type II) or stable elevation (Type III). See **Fig. 3c** for sample traces for each type. Bars represent mean ± standard error mean for each target membrane, as indicated. No statistically significant difference between target membranes was found.

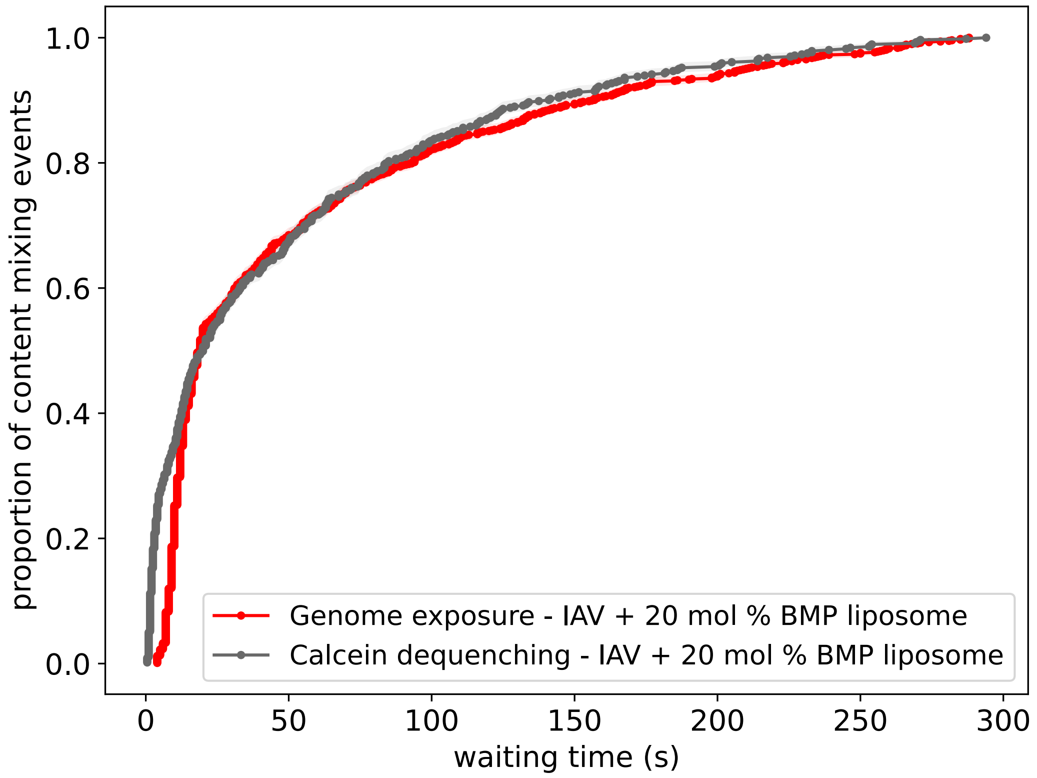


**Figure S2. Comparison of content mixing kinetics between different fluorophores.** The cumulative distribution functions (CDFs) of event waiting times for IAV bound to 20 mol % BMP liposomes loaded with a nucleic acid-binding dye (DiYO-1), described previously by Mannsverk, Villamil Giraldo, and Kasson (2022) or Calcein, as described here. The genome exposure CDF is replotted from the data in Mannsverk, Villamil Giraldo, and Kasson (2022). Note that the genome exposure CDF is right-shifted by a few seconds due to the difference in how time 0 was determined: For the genome exposure assay, time 0 = when the buffer exchange is initiated, while for the calcein dequenching assay time 0 = when the background calcein signal drops due to the pH sensitivity of calcein. The latter is thought to be a more precise measure of when the viral particle is exposed to the low pH buffer.

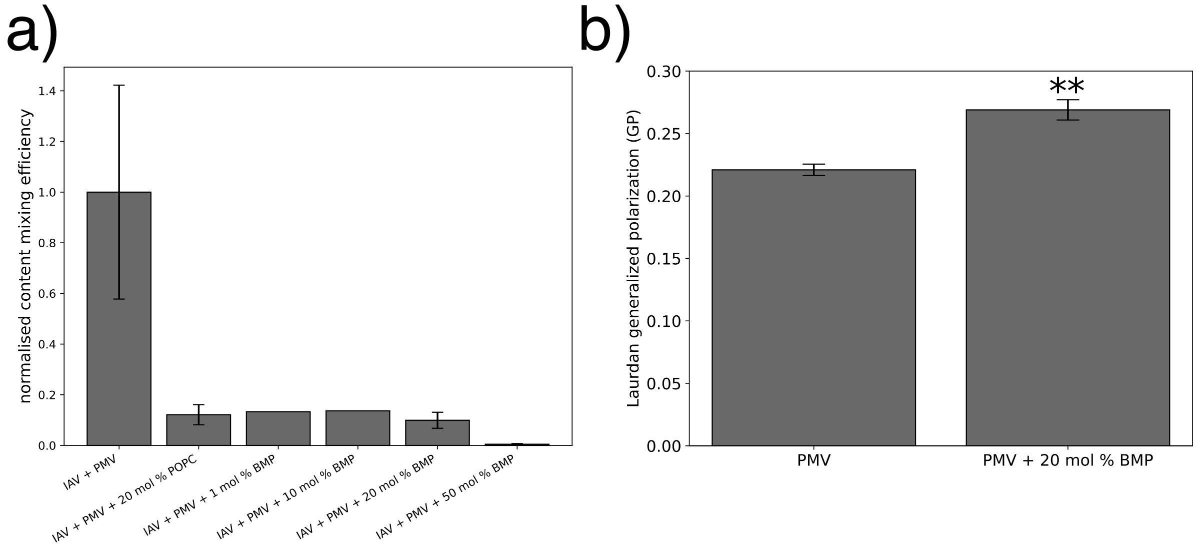


**Figure S3. Content mixing efficiency and Laurdan generalized polarization of PMVs directly supplemented with exogenous lipids. a**) Content mixing efficiency is plotted between IAV and PMVs supplemented with BMP or POPC. For these experiments, the exogenous lipid was dissolved in chloroform followed by shaking at 800 rpm at room temperature for 30 mintues to promote lipid incorporation. The amount of lipid to add to the PMV sample was estimated (see **Supplementary methods item 1**) and the volume of solvent added to the PMV sample was less than 1 % of the total sample volume. Bars show mean mixing efficiency, calculated content mixing events / HA-positive particles in each field of view. Error bars represent standard error and are plotted for samples with at least 3 repeats. Values are normalised to the content mixing efficiency of untreated PMVs. **b**) Laurdan generalized polarization (GP) is plotted for PMVs supplemented with BMP directly prior to labelling with C-Laurdan. A one-way ANOVA and Tukey HSD post-hoc test was carried out to determine significant difference between groups. ** signifies a p-value < 0.01. See **Fig. 5b** for more information on Laurdan GP.

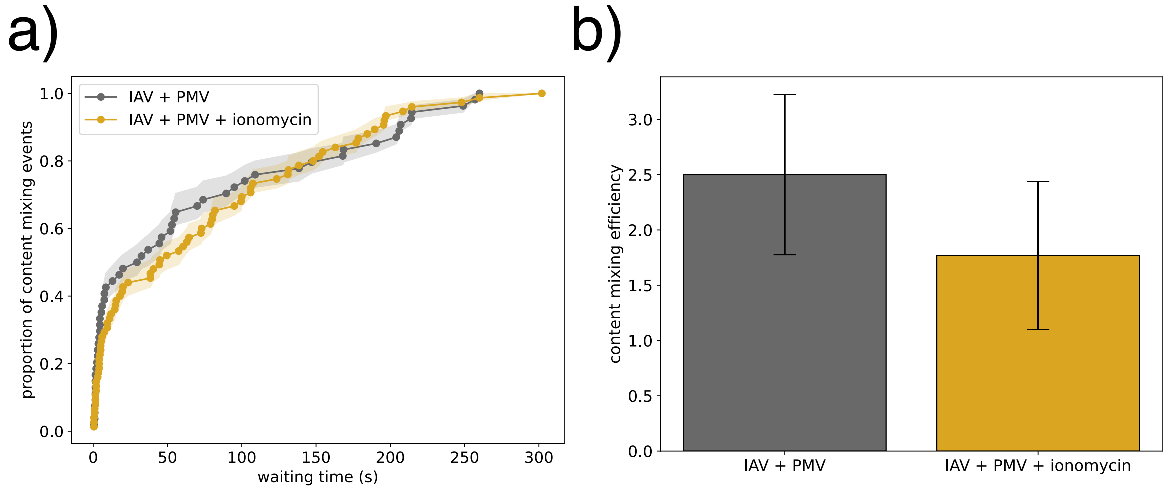


**Figure S4. Content mixing kinetics and efficiency of IAV with lipid-scrambled PMVs.** PMVs bound to a flow cell channel were pretreated with 10 μM ionomycin for 30 min at 37°C, before the ionomycin was washed out and IAV introduced. **a**) Kinetics are plotted as normalized CDFs of single-event waiting times. No statistically significant difference between the CDFs was found by a KS test (p-value = 0.58) or bootstrapped rank sum test over flow cells (p-value = 0.98). **b**) Content mixing efficiency is plotted, calculated as total events recorded / TR-labeled IAV particles detected in the same field of view. Bars show mean ± standard error. No significant difference between group was found using a two-sample t-test (p-value = 0.50)

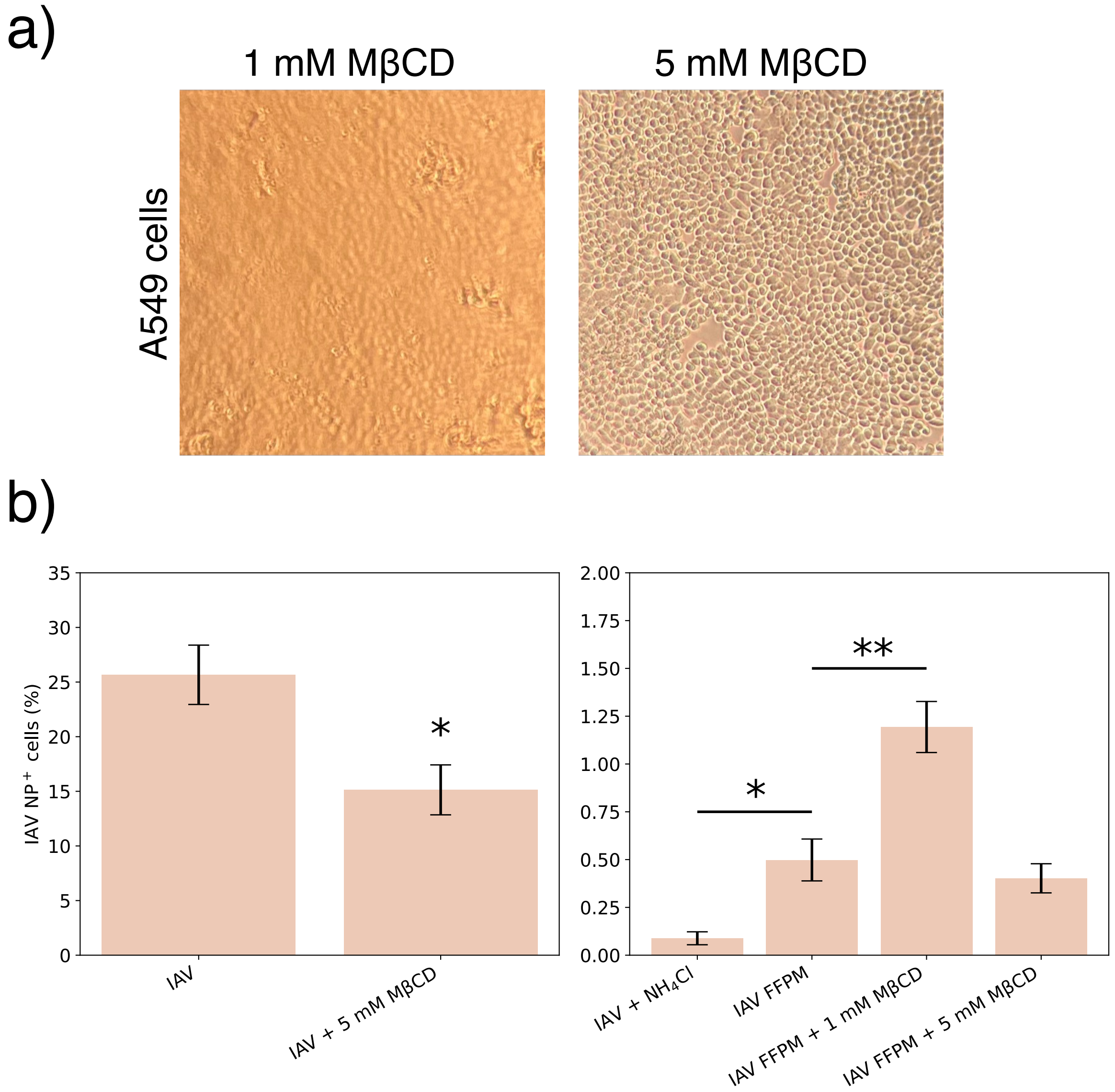


**Figure S5. IAV FFPM in A549 cells pre-treated with MβCD. a**) bright-field images of an A549 monolayer after treatment with 1 mM (left image) or 5 mM (right image) MβCD for 30 min at 37°C. Cells in the right image show loss of cell-cell adherence after MβCD treatment. However, the cells remained firmly attached to the bottom of the well throughout the FFPM infection experiment. **b**) Quantification of IAV NP-positive cells 5 hours post-infection. NP positivity was defined as a mean nuclear NP intensity above uninfected cells, and the percent NP-positive nuclei was calculated in each sample as in **Fig. 6b**. A one-way ANOVA and Tukey HSD post-hoc test was used to determine statistically significant differences between groups, with significantly different groups labelled on the plot. Bar plot shows mean of each group ± standard error mean from three repeats. * and ** indicate p-values < 0.05 and < 0.01, respectively. Note different y-axis range on left and right panel. However, experiments plotted in both panels were carried out in parallel and are therefore comparable.
